# Circadian clock-gated cell renewal controls time-dependent changes in taste sensitivity

**DOI:** 10.1101/2023.03.09.531858

**Authors:** Toru Matsu-ura, Kaoru Matsuura

## Abstract

Circadian regulation of the cell cycle progression produces a diurnal supply of newborn cells to replace the ones that were lost in the organs and tissues. Here we analyzed time-dependent changes in the cell types in the mouse tongue epithelium. We observed circadian regulated alternate oscillations of the stem/progenitor cell maker genes and the differentiated cell marker genes expressions in mouse tongue epithelial organoids. The cell cycle progression was regulated time-dependent manner in the tongue organoids and mice tongue. Single-cell RNA sequencing revealed time-dependent population changes of the stem/progenitor cells and the differentiated cells in mice tongues. Remarkably, we observed time-dependent type II taste cell population changes, resulting in time-dependent taste sensitivity changes. We also found the same population changes in mice intestines and uteruses, suggesting the contributions of the diurnal supply of newborn cells to the time-dependent physiological controls in the broad types of organs and tissues.

## Introduction

The circadian clock and the cell cycle are biological oscillators whose coupling is observed from cyanobacteria to humans (Bjarnason et al., 1999; Hong et al., 2014; Yang et al., 2010). In mammals, daily rhythms of the cell divisions were observed in various types of cells from cultured cancer cells to adult organs and tissues. The adult stem cells and the progenitor cells are major proliferating cells in the adult tissues. In the rodents’ skin and hair follicles, the progenitor cells are the major proliferating cells, and the circadian clock gates the cell cycle in epidermal stem and progenitor cells, which controls the timing of S-phase at around the late afternoon to provide protection against DNA damage from UV irradiation (Geyfman et al., 2012; Plikus et al., 2013). Although the timing of cell divisions in the day is varied in the types of tissues, the dairy cell division cycles were observed though out the body including the skin, brain (Guzman-Marin et al., 2007), cornea, gastro-intestinal tract (Sheving, 2000, Gastroenterology), bone marrow, blood cells (Smaaland et al., 2002), circulating hematopoietic stem and progenitor cells (Mendez-Ferrer et al., 2008), and oral epithelium (Bjarnason et al., 1999). The circadian regulation of the cell cycle progression is known to be achieved by the circadian regulation of cell cycle components including checkpoint kinases and cyclin-dependent kinases (Grechez-Cassiau et al., 2008; Kowalska et al., 2013; Matsu-Ura et al., 2016; Matsuo et al., 2003).

The clock-gated cell cycle progression is also found in the tongue. Kummermehr’s group reported the circadian rhythms of the mouse tongue epithelial proliferation whose DNA synthesis (S)-phase is in the night and the mitosis (M)-phase is in the morning (Dorr and Kummermehr, 1991; Moses and Kummermehr, 1986). Aihara *et al*. also reported 24 h rhythm of the cell cycle progression using a cell cycle sensor, FUCCI2, expressing taste bud organoids (TBOs) (Aihara et al., 2015). Passage through the cell cycle is coordinated by cyclin proteins, cyclin-dependent kinases (cdks), and checkpoint proteins. The circadian variation in the nuclear expressions of those proteins was also reported with distinct peak times of each protein to achieve time-dependent cell cycle progression (Bjarnason et al., 1999).

Including the cell cycle progression, the circadian clock regulates the variety of cellular and tissue physiological events. In *Drosophila*, synchronous rhythms in physiological and behavioral responses to attractive and aversive tastants are under clock control (Chatterjee and Hardin, 2010). The same circadian variation of the taste recognition threshold is also found in humans. Two groups separately reported sensitization to the salty taste in the afternoon (Fujimura et al., 1990) and the sweet taste in the morning (Nakamura et al., 2008).

Here, our work connecting the circadian-gated cell cycle progression and the diurnal variation of taste sensitivity represents the importance of synchronized cell cycle progressions to the functions of tissues and organs. Using single-cell RNA sequencing and flow cytometry, we demonstrated that synchronized cell cycle progressions produce time-dependent population changes of the stem/progenitor cells and the differentiated cells in the mice tongues. Remarkably, we observed time-dependent type II taste cell population changes, which resulted in time-dependent changes in taste sensitivity. In the mouse tongue epithelium, not only the cell cycle but also apoptosis of the cells are under clock-controlled, suggesting both the production and the elimination of the functional epithelial cells regulate the population changes of the tissue. We observed the same population changes in the intestinal and uterine epithelium. Together, these results demonstrate that the circadian-clock-cell cycle coupling can work in multiple epithelial tissues in the body to control functional variation in the day.

## Results

### Circadian regulation of stem/progenitor and differentiated cell marker genes in TBOs

We are interested in the gene expressions to the functional variations of the epithelial tissues of mammals. We developed TBOs from mouse tongue epithelium and checked the circadian regulations of gene expressions (Aihara et al., 2015). We found expression levels of core clock genes *Bmal1* and *Per2* in the TBOs have 24 h oscillatory rhythms (Figures 1a and b). The oscillatory gene expressions are also found in stem cell marker genes including *Lgr6*, *Axin2*, and *Bmi1* (Figure 1c-e). The peaks of gene expression of these three genes appeared same time to form synchronized oscillations. We also detected synchronized oscillations of differentiated taste cell marker genes *Krt8*, *Gnat3*, and *NTPDase2* (Figure 1f-h). Interestingly, peaks of the stem cell marker genes and the differentiated taste cell marker genes appear reciprocally, suggesting the formation of both types of cells at different times of the day. The oscillatory expressions of those genes were not observed in TBOs with *Bmal1* knockdown with the shRNA (Figure S1a) (Matsu-Ura et al., 2016).

**Figure 1.**
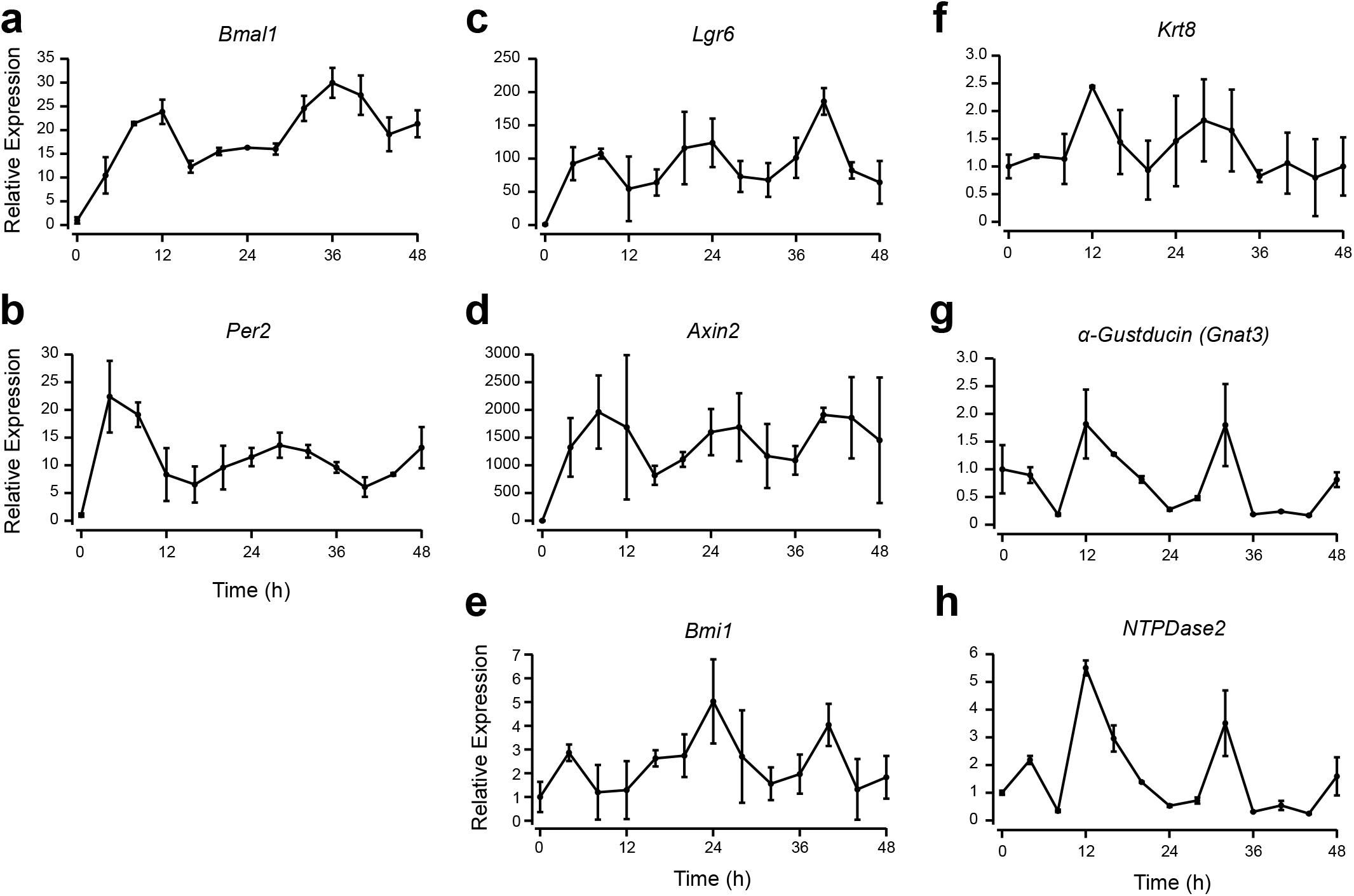
Oscillatory expression of stem/progenitor and differentiated cell marker genes in TBOs. (**a-b**) mRNA expression of core clock genes, *Bmal1* (**a**) and *Per2* (**b**). (**c-e**) mRNA expression of adult stem cell marker genes, *Lgr6* (**c**), *Axin2* (**d**), and *Bmi1* (**e**). (**f-h**) mRNA expression of differentiated cell marker genes, *Krt8* (**f**), *α-Gustducin* (*Gnat3*) (**d**), and *NTPDase2* (**e**).

### Diurnal variation of cellular populations in the mouse tongue epithelium

To understand the variations in each cell type of epithelial tissue, we performed single-cell RNA sequencing (scRNAseq) of mouse tongue epithelial cells collected at Zeitgeber time (ZT) 0, the time of lights on, and 12, the time of lights off. We eliminated dead or dying cells by the expression of mitochondria genes (Stuart et al., 2019) and analyzed 1405 cells of ZT0 and 2352 cells of ZT12 (Figure 2a). By expressions of stem/progenitor and tongue epithelial cell marker genes (Figure S2a-c), we classified 11 identified clusters by principal component analysis into four different subtypes; stem cells, progenitor cells, filiform papillae cells, and mucosal non-keratinized epithelial cells (Figure 2a). Figure 2b shows the comparisons of populations of each cell type. The progenitor cells are the largest population at ZT0 (39.9% at ZT0 and 31.5% at ZT12, Figure 2b), and the mucosal non-keratinized epithelial cells are the largest population at ZT12 (36.0% at ZT0 and 48.3% at ZT12, Figure 2b). These results suggest that the tongue epithelium has the diurnal variation of the cellular populations. The taste cell marker gene-expressing cells are distributed throughout the clusters and did not form specific clusters in the UMAP graph (Figure S2d). Thus, we counted the cells which express type I, II, III, and pan taste cell marker genes to quantify the diurnal population changes of each type of taste cell (Table S1). As shown in Figure 2c, the population of cells with expression of type II taste cell marker genes is greater in ZT0 compared to that in ZT12. Using TBOs, we next measured the taste cell populations by flow cytometer. The circadian clock of the TBOs was synchronized by the serum shock, and then the taste cells were analyzed. As same as the result obtained with scRNAseq, we found the cells with type II taste cell marker GNAT3 expressing populations are greater at 12 h after the serum shock (SS12) than that at SS24 (Figure 2d and S2e). We also obtained the same data using cells obtained from mouse tongue epithelium (Figure 2e and S2f).

**Figure 2.**
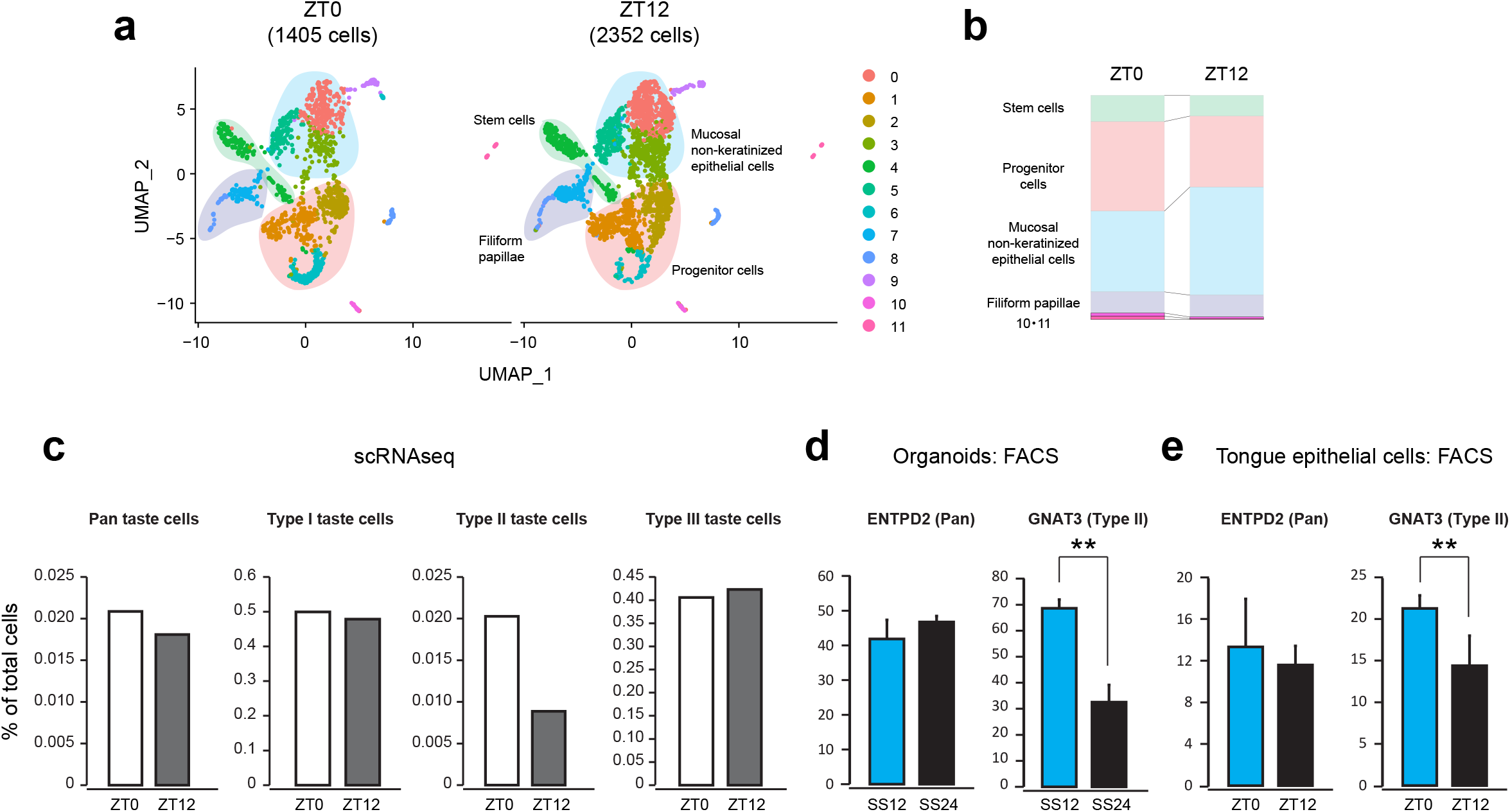
Cellular population changes in the mouse tongue epithelium at different circadian time points. (**a-c**) The results of scRNAseq of mouse tongue epithelium. (**a**) Uniform manifold approximation projection (UMAP) plot depicting clustering of the single tongue epithelial cells using the scRNAseq data with annotations of four cell types, stem cells (green background), progenitor cells (pink background), filiform papillae cells (purple background), and mucosal non-keratinized epithelial cells (blue background). The left panel shows data from the ZT0 mouse, and the right panel shows data from the ZT12 mouse. (**b**) Stacked bar graphs show differences in the tongue epithelial cell populations at the ZT0 and the ZT12. (**c**) Differences in taste cell populations at the ZT0 and the ZT12. Cells with pan taste, type I taste, type II taste, and type III taste marker gene expression were counted. (**d-e**) Circadian-dependent taste cell population changes were detected with flow cytometry experiments. (**d**) Differences in taste cell populations of TBOs at the time points of SS12 and SS24. (**e**) Differences in taste cell populations of mouse tongue epithelium at the time points of ZT0 and ZT12. Bars and errors correspond to the average values and the SDs, respectively. \*\**p* < 0.01, student’s *t*-test.

### Circadian clock regulated cell cycle progressions in the mouse tongue epithelium

We performed gene enrichment analysis of the cell cycle relating genes in the scRNAseq data and defined the cells in the gap 1 (G1), S, and gap 2 (G2)-M phases (Hao et al., 2021) to find differences of the cell cycle phases at the two-time points (Figure 3a). We found that the population of the cells in the G1 phase is greater at ZT12 and that in the G2-M phase is greater at ZT0 (Figure 3b), suggesting the clock-gated cell cycle regulation in mouse tongue epithelium. To confirm the clock regulation of the cell cycle, we developed reporter TBOs by expressing a bioluminescent circadian reporter Bmal1-eLuc (Figure 3c) and a bioluminescent cell cycle reporter GreenLuc-hGeminin (Figure 3d) (Matsu-ura 2016). We found 24 h rhythms of bioluminescent signals obtained from both Bmal1-eLuc and GreenLuc-hGeminin (left panels of Figure 3c and d). As reported in our previous publication, the circadian gene expressions and the regulations in the intestinal epithelium are cell-type specific and diverse (Matsuura 2016). The circadian oscillations in the intestinal stem cells are less robust, and the cell cycle of the intestinal stem cells is regulated through the circadian-regulated WNT secretion from Paneth cells. We tested a stem cell-rich culture medium (Matsuura 2016) to TBOs and found upregulation of a stem cell marker gene *Lgr6* and reduced expression of differentiated cell marker genes *Gnat3*, *NTPDase2*, and *K8* (Figure S3a). In the stem cell-rich condition, both Bmal1-eLuc and GreenLuc-hGeminin signals show less robust oscillations (right panel of Figure 3c and the middle panel of Figure 3d). We also found less robust GreenLuc-hGeminin oscillations in a core clock gene, *Bmal1*, knockdown TBOs, showing the circadian regulation of the cell cycle progression in TBOs (right panel of Figure 3d).

**Figure 3.**
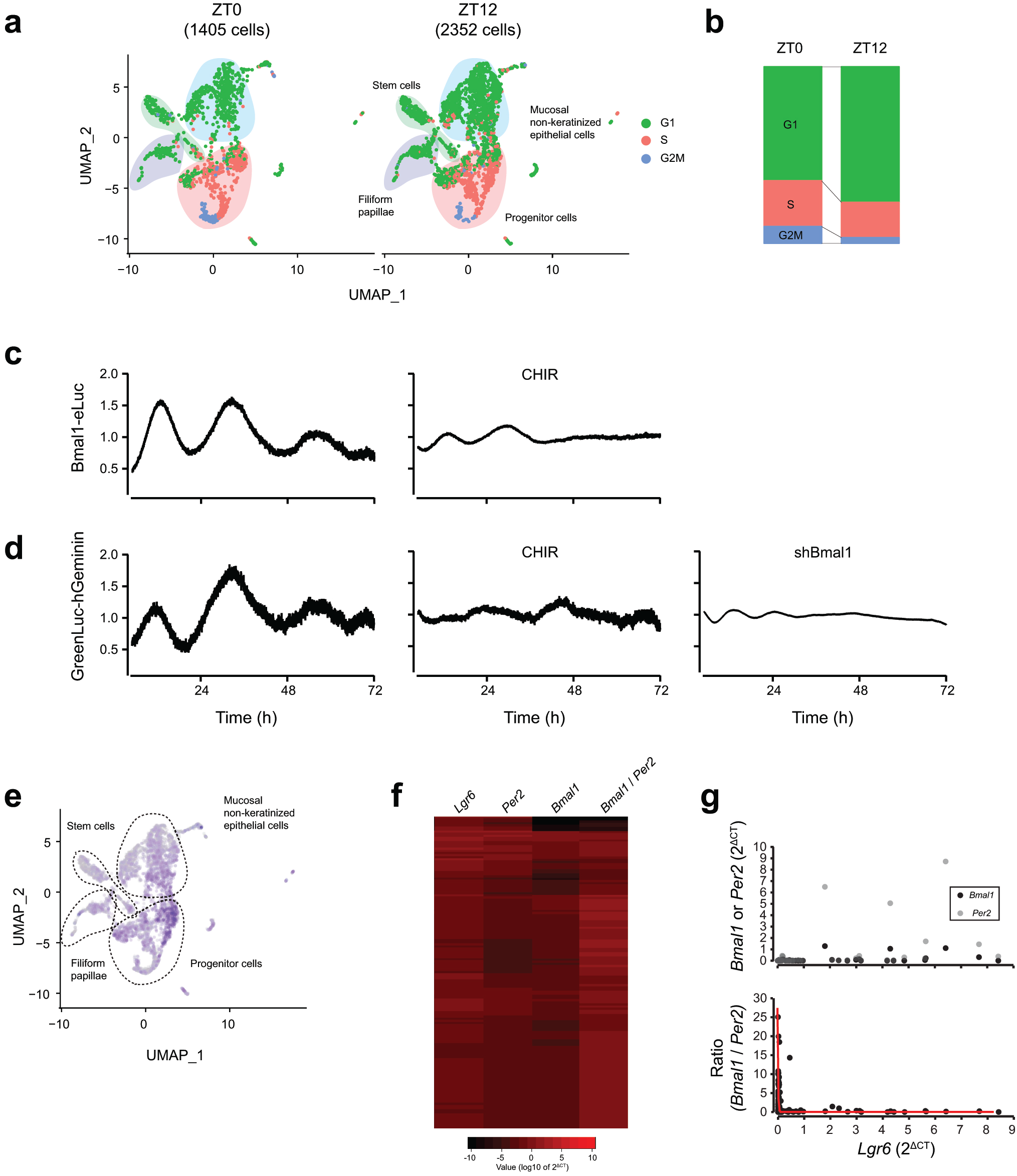
Circadian-regulated cell cycle progressions in the mouse tongue epithelium. (**a**) UMAP plot with the phase information of the cell cycle. Cells in G1, S, and G2M phases are represented by green, red, and blue dots, respectively. (**b**) Stacked bar graphs show differences in the cell cycle phases of tongue epithelial cell populations at the ZT0 and the ZT12. (**c**) Time course changes of a circadian reporter Bmal1-eLuc in control TBOs (left panel) and a WNT activator CHIR-treated TBOs (right panel). (**d**) Time course changes of a cell cycle reporter GreenLuc-hGeminin in control TBOs (left panel), CHIR-treated TBOs (middle panel), and Bmal1 knock-down TBOs (right panel). (**e**) The superimposed UMAP plot of the 13 core-clock gene expression maps (Figure S3). (**f-g**) Single-cell qPCR (scqPCR) analysis of TBOs at SS12. (**f**) A heat map of the scrPCR results. (**g**) The graphs show relationships of expression levels of *Lgr6* vs *Bmal1* or *Per2*(upper panel) and the expression level of *Lgr6* vs *Bmal1* and *Per2* expression ratio (lower panel).

The heterogeneous expressions of circadian clock genes are also found in the scRNAseq data from mouse tongue epithelium. We created feature maps of core clock genes expressing in the daytime (Figure S3b), the nighttime (Figure S3c), and the superimposed image of the maps of 13 core-clock genes (Figure S3d and Figure 3e). The superimposed image shows the core clock genes express more in the population of progenitor cells and less in the population of stem cells (Figure 3e). To further confirm heterogeneous circadian regulations in tongue epithelium, we performed single-cell real-time PCR (scRT-PCR) of TBOs at SS12, the time of high *Bmal1* expression (Figure 3d), to compare the clock gene expressions and the stem cell marker gene, *Lgr6*, expression. As we expected, expressions of core clock genes, *Per2* and *Bmal1*, are heterogeneous, and we found the cell population with low expression of *Baml1* (Figure 3f). We found an exponential decay of *Bmal1/Per2* expression ratio against *Lgr6* expression level (Figure 3g). These data suggest that the regulation of the circadian clock can be varied in the cell types and robust circadian-gated cell cycle regulation in the proliferating progenitor cells in the tongue epithelium.

### Diurnal variation of the cell death rate and the cell populations in tongue, intestinal, and uterine epithelium

We found the population changes of stem/differentiated cells and differentiated cell types in the mouse tongue epithelium. The daily supply of the newer divided cells is a major candidate to make time-dependent cellular population changes in tongue epithelium. On the other hand, eliminating old cells also has the potential to make time-dependent population changes. Using an active-CASPASE3 antibody, we counted apoptotic cells in the tongue epithelium and found more apoptotic cells in ZT0 than in ZT12 (Figure 4a, b, and S4a). We also found more apoptotic cells in ZT0 in mouse intestinal and uterine epithelium cells (Figure 4c-d and S4a-b). In the intestinal cell layer, we found more programmed cell death in the villi where the differentiated cells locate compared to that in the CRYPTS where stem/progenitor cells locate (Figure 4c). KRT18 and MUC2 are the markers of tuft and goblet cells in the intestine, respectively (Hofer and Drenckhahn, 1996; Kim and Ho, 2010). We found more KRT18 positive cells at ZT0, but the time-dependent population change was not detected for MUC2 positive cells (middle and right panels of Figure 4c). We also detected more stem/progenitor cells at ZT12 in the uterine epithelium (Figure 4d and S4b).

**Figure 4.**
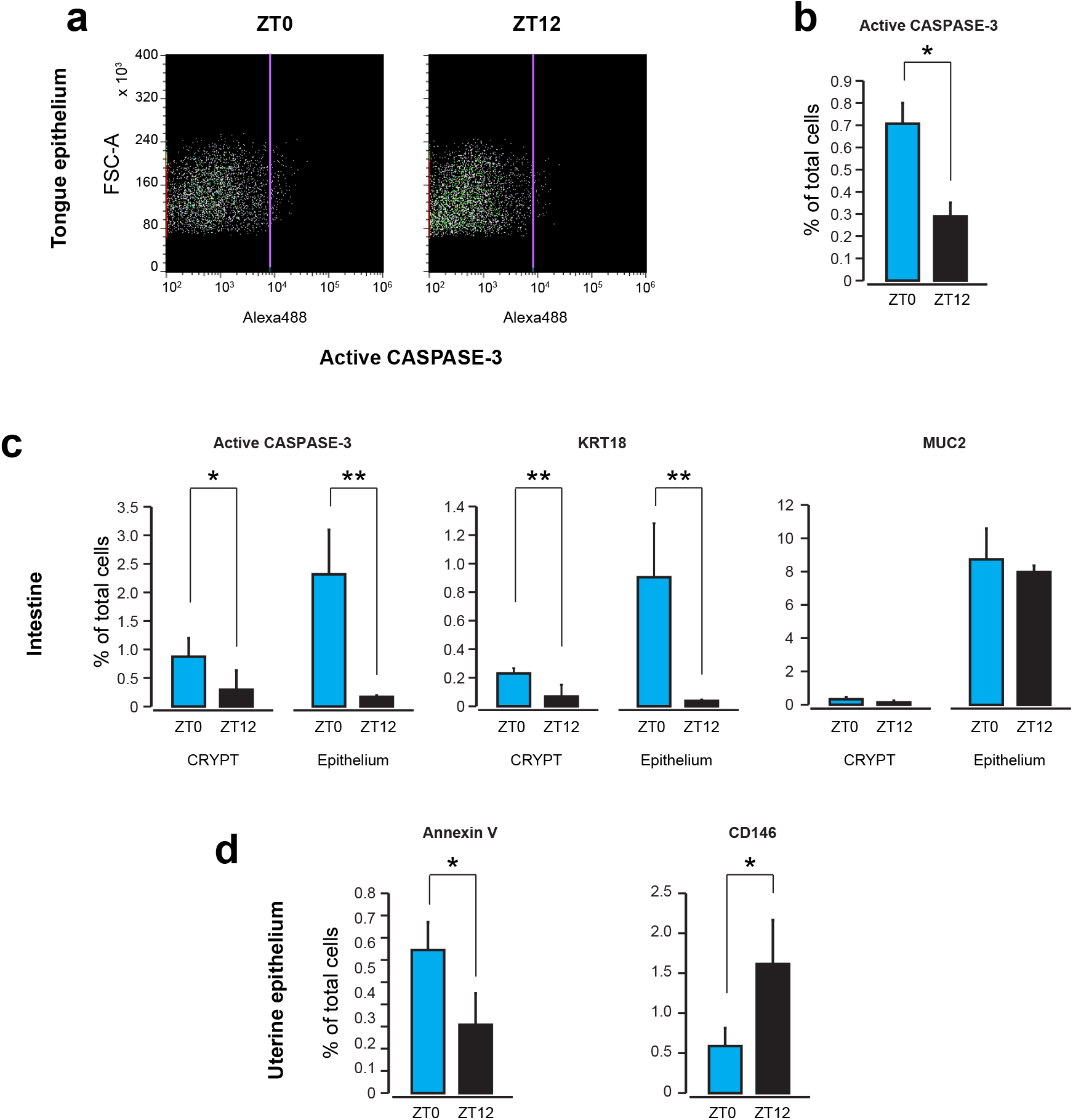
Diurnal variation of the cell populations and the cell death rate in tongue, intestinal, and uterine epithelium. (**a-b**) Flow cytometric analysis of apoptotic cells in the mice tongue epithelium. Cell scatter graphs (**a**) and a bar graph (**b**) are shown. (**c**) Flow cytometric analysis of apoptotic cells (left panel), tuft cells (KRT18, middle panel), and goblet cells (MUC2, right panel) in the mice intestinal epithelium. (**d**) Flow cytometric analysis of apoptotic cells (Annexin V, left panel) and stem cells (CD145, right panel) in the mice uterine epithelium. Bars and errors correspond to the average values and the SDs, respectively. \**p* < 0.05, \*\**p* < 0.01, student’s *t*-test.

### Diurnal variation of the cell populations regulates taste sensing in tongue epithelium

To find out the functional consequences of the cell population variation in a day, we studied the recognition threshold of tastants in mice. Wild-type (WT) mice were housed in two isolators whose light/dark cycles have a 12 h time difference. Those mice were subjected to two water bottles; one is pure water and the other is water with tastants at ZT0 or ZT12. We tested three different concentrations of each tastant to find the threshold concentrations. We first tested tastants to type II taste cells which sense bitter, sweet, and umami. Denatonium benzoate (DB) is a bitter tastant, and the mean preference ratios of DB at three concentrations are all less than 0.5, meaning that the mice avoid drinking DB-containing water (left panel of Figure 5a). We found a significant difference in the preference for DB-containing water at the DB concentration of 0.2 mM. Mice avoid drinking 0.2 mM DB water more at ZT0 compared to that at ZT12. The same sensitivity differences were also obtained with sweet and umami tastants. Mice are more sensitive to sucrose or monosodium glutamate (MSG) containing water at ZT0 (middle and right panels of Figure 5a). On the contrary, we did not detect any significant differences in the sensitivity to salty and sour tastants, which are detected by type I and III taste cells, at the two-time points (Figure S5a). We next tested *Lgr6*^*CreERT*/+^/*R26*^*DTA*/+^ mice for the recognition threshold of tastants. Tamoxifen (TX) solution was injected into mice three times every other day to kill *Lgr6*-expressing tongue epithelial stem cells by the expression of diphtheria toxin (DTA) (Huang et al., 2021), which resulted to stop the daily refilling of newly produced cells. The tamoxifen-treated mice were tested for the tastants after three days of the last administration. In contrast to WT mice, *Lgr6*^*CreERT*/+^/*R26*^*DTA*/+^ mice did not show time-dependent differences in threshold changes to the type II taste cell tastants; DB, sucrose, and MSG (Figure 5b).

**Figure 5.**
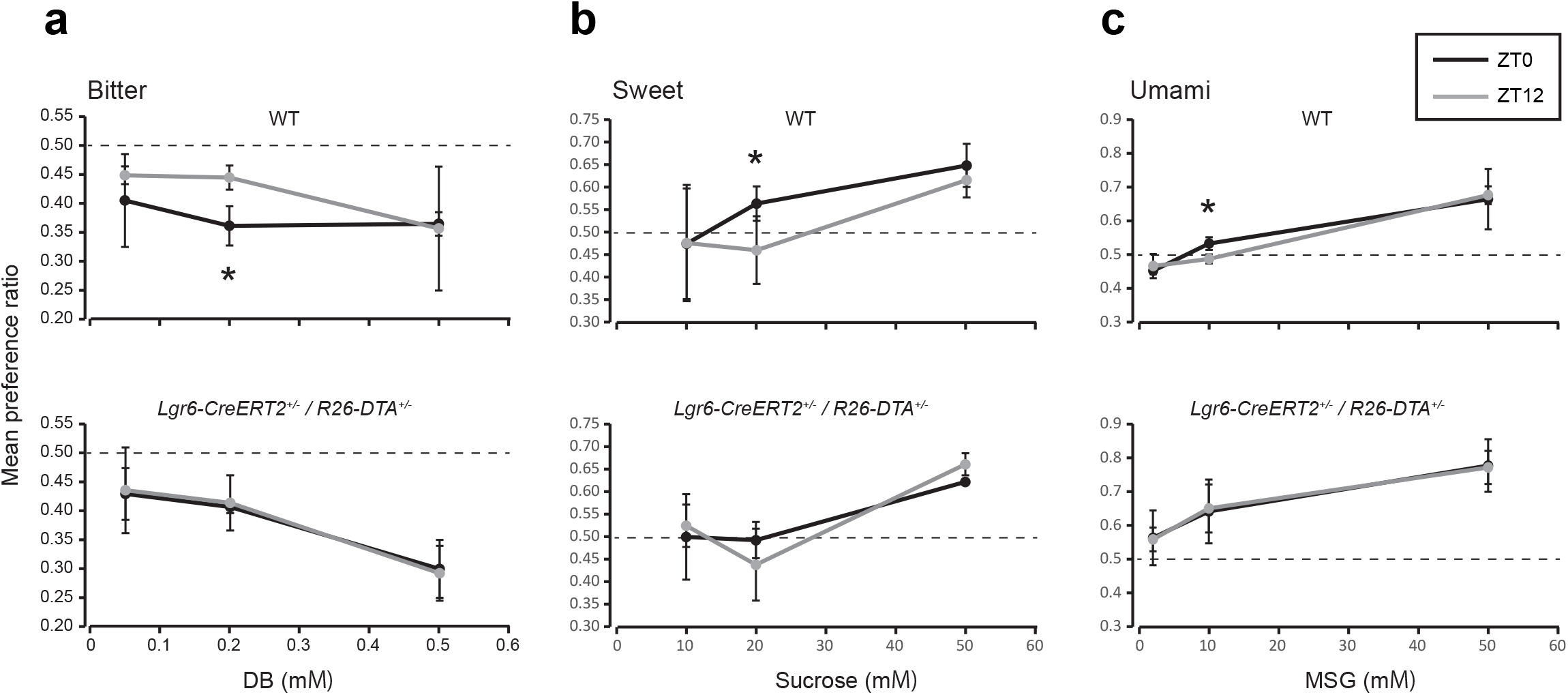
Diurnal variation of the taste sensing in tongue epithelium. Mean taste preference ratios of ZT0 and ZT12 mice using 6-hour two-bottle (tastant versus distilled water) preference tests. Behavioral responses of WT (upper panels) and *Lgr6*^*CreERT*/+^/*R26*^*DTA*+^ mice (lower panels) to bitter (**a**, DB: 0.05, 0.2, and 0.5 mM), sweet (**b**, Sucrose: 10, 20, and 50 mM), and umami (c, MSG: 2, 10, and 50 mM). The values and errors correspond to the average values and the SDs, respectively. \**p* < 0.05, student’s *t*-test.

## Discussion

Circadian-gated cell divisions provide daily refills of newly produced cells to replace old or damaged cells in our body. The newly produced cells are dispensable to maintain the function of tissues and organs. The meaning of the circadian-gated cell divisions is well understood in skin and hair follicles. These tissues are located on the surface of organisms and face the threat of DNA damage by UV irradiation. Thus, clock gates DNA synthase around the late afternoon to avoid UV light in the sunlight (Geyfman et al., 2012; Plikus et al., 2013). On the contrary, tissues and organs inside of our body are not directly irradiated by UV light. Thus, the diurnal variation inside of our body may not have a significant meaning to the survival of the cells. In fact, cells in some tissues are known to synthesize DNA in the daytime. Our findings of the diurnal variation of cellular population changes in tongue epithelium, together with time-dependent changes in taste sensitivity threshold, add the first evidence of population changes of cell type and functional changes in the tissue, which are produced by circadian-gated cell cycle progression. In this report, we demonstrated alternate peak appearances of circadian oscillated stem cell markers and differentiated marker genes in TBOs (Figures 1 and S1), which should be the result of circadian-gated cell cycle progression (Figure 2d). The results suggest that progenitor cells are produced at a specific time of the day and differentiate at other times depending on the types of differentiated cell types. The same time-dependent differentiation control is also reported in an embryonic stem cell differentiation system which shows a daily burst of differentiation and production of adipocytes by the circadian control of PPARG2 gene expression (Zhang et al., 2022). Our scRNAseq of mouse tongue epithelium also has the diurnal variation of differentiated cells, showing the daily burst of differentiation *in vivo* (Figure 2b). Importantly, the population change is not shown in all types of differentiated cells. Our flow cytometry analysis shows population changes in type II taste cells but not in total taste cells (Figure 2e). The same result was observed in the intestinal epithelium. We observed the time-dependent population change in tuft cells but not in goblet cells (Figure 4c).

In our scRNAseq data, we observed cell-to-cell variations in the expression levels of circadian relating genes (Figure 2e). The circadian relating gene expressions are high in progenitor cells and low in stem cells (Figure 2e). Yagita’s group continuously reported circadian development during the differentiation of pluripotent stem cells. Pluripotent stem cells lack functional circadian transcriptional/translational feedback (Yagita et al., 2010) because of the absence of CLOCK protein, and the posttranscriptional regulation of CLOCK plays a role in setting the timing for the emergence of the circadian clock oscillation during differentiation of pluripotent stem cells and mouse embryos (Umemura et al., 2017). Less robust circadian oscillations in adult stem cells are also reported in mouse intestinal organoids, and divisions of the intestinal epithelial stem cells are under the control of the circadian clock in Paneth cells through the secretion of WNT (Matsu-Ura et al., 2016). In tongue epithelium, we find higher circadian gene expressions in the progenitor cells which are the major proliferating cells in the tissue (Figure 2e), conforming to the observed circadian-gated cell cycle progression (Figure 2d). Furthermore, our scRNAseq and scqPCR data show single-cell level circadian variability (Figures 2e-g) which can be explained by the stochasticity of gene expression levels and clonal heterogeneity with experiments with U2OS cells (Nikhil et al., 2020).

We find not all taste cells but only type II taste cells have population changes in a day, which resulted in the differences in taste sensing of bitter, sweet, and umami tastants (Figure 5 a). The tendency of population change can be dependent on the lifetime of each type of cell in the tissue (Seim 2017). One interesting phenomenon which can regulate cellular lifetime was reported by Bao et al. They showed reduced cell death on the villus of the TRα knockin mutant intestine because of cell proliferation decreases in the crypt (Bao et al., 2019), which can be explained by cell death stimulation with “push” pressure and accompanied changes in cell-ECM interaction (Bao et al., 2020). In our system, we slowed the cell proliferation by DTA expression in *Lgr6*+ taste bud epithelial stem cells, which can elongate the lifetime of taste cells by reducing the “push” pressure. Although the diurnal taste sensing difference should be the result of diurnal type II taste cell population change, we do not know the meaning of the taste sense difference in the survival of animals.

Nakamura *et al*. reported diurnal variation of human sweet taste recognition thresholds. They found 21-30 years old human subjects have higher sweet sensitivity in the morning (Nakamura et al., 2008) as same as our mouse experiment showing that higher sweet sensitivity at ZT0 compared to that at ZT12 (Figure 5a). For human health in developed countries, it is possible to reduce the amount of sugar in the morning since humans have higher sensitivity in the morning. Reduced proliferation of taste cells in *Lgr6*^*CreERT*/+^/*R26*^*DTA*/+^ mice is a part of the reproduction of cellular senescence in aged animals with a slow turnover of taste cells. Taste threshold changes in *Lgr6*^*CreERT*/+^/*R26*^*DTA*/+^ mice suggest that reduced circadian-regulated cellular population changes are one of the reasons for taste-sensing changes in aged persons (Wysokinski et al., 2015).

In conclusion, we uncovered diurnal cellular population changes in type II taste cells, which resulted in diurnal variation of the taste-sensing threshold of bitter, sweet, and umami tastants. Cell divisions of most cells in our body are under circadian control. As we find the same cellular population changes of KRT18-positive tuft cells in the intestine, these population changes can be observed in more tissues. Thus, some parts of the circadian variation of tissue functions can be regulated by cellular population changes.

## Experimental procedures

### Animals

Experiments used C57BL/6J (SLC), *Lgr6*^*CreERT*/+^ (Jackson Laboratory), and *R26*^*DTA*/+^ (Jackson Laboratory) mice. All mice were housed in the animal facility at Kansai Medical University (KMU). All animal experiments were approved by KMU welfare committee.

### Preparation of TBOs

Using the methods of Aihara *et al*. (Aihara et al., 2015), we prepared taste bud organoids (TBOs) by isolating fresh tongue epithelium from mice. Dispase solution (Corning: 354235) was injected into isolated tongues from sacrificed animals, and incubated at 37 □ for 15 minutes. Then, the epithelium sheets were removed, minced by scissors, and digested by TrypLE Express enzyme mix (ThermoFisher: 12604021). The dissociated cells were isolated by filtration to remove debris with 70 μm cell strainer (Falcon: 352350) and centrifuge. The isolated cell were embedded in extracellular matrix gel (ThermoFisher: A1413202) droplets and cultured with Advanced DMEM/F12 (ThermoFisher: 12634010) supplemented with 50% WNT conditioned medium, 10% R-spondin conditioned medium, 10% Noggin conditioned medium, B27 (ThermoFisher: A1486701), N2 (ThermoFisher: 17502001), EGF (Peprotech: AF-100-15), Hepes pH7.4, L-Glutamine, Penicillin, and Streptomycin.

### Lentivirus induction

TBOs were treated by TrypLE Express and dissociated by pipetting. Dissociated cells were suspended in the culture medium, 10 μM Y-27632 (ROCK inhibitor, Sigma-Aldrich: Y0503), and lentivirus. Bmal1-eLuc and GreenLuc-hGeminin expressing TBOs and Bmal1-KD TBOs were selected by puromycin (1 μg/ml).

### Bioluminescence recording

TBOs were plated on 35 mm plastic dish with the culture medium and 200 μM Beetle Luciferin K+ salt (Nacalai: 20028-24). The plates were placed in a CO_2_ incubator with a luminometer (Hamamatsu: H9319-02) for real-time periodic quantification of Bmal1-eLuc and GreenLuc-hGeminin abundance by bioluminescence recording.

### qPCR

TBOs were plated into 24-well plates at the start of the experiment. Organoids for each time point were plated into a separate plate to limit manipulation or exposure to possible resetting cues. End of serum shock is indicated by circadian time 0 to compare with bioluminescent recordings performed in parallel. Total RNA was harvested every 4 h over a 48-h period using 0.25 ml/well of Trisole Reagent (Sigma-aldrich: T9424). For reverse transcription reactions, 0.2-1 μg of total RNA was used with Superscript III (ThermoFisher: 12574018) according to manufacturer’s instructions. qPCR was performed with SYBR Green Master Mix (Qiagen, Germantown, MD, USA) using primers. qPCR results were detected by using the StepOne System (ThermoFisher).

### Single cell RNA sequencing (scRNAseq)

Mice were sacrificed at ZT0 and ZT12. Tongue epithelial cells were isolated as the method of preparation of TBOs. Dead cells were stained with 7-AAD (ThermoFisher: A1310) and removed with cell sorter (SONY: SH800S). The cells were then processed using the Chromium Controller (10×Genomics, Pleasanton, CA), Chromium Next GEM Single Cell 3’ GEM, Library and Gel Bead Kit v3 (10×Genomics; PN-1000128), and MGIEasy Universal Library Conversion Kit (App-A) (MGI, Shenzhen, China) following the manufacturer’s instructions. The library was sequenced using DNBSEQ-G400 (MGI).

### Downstream analysis of 3’-scRNA-seq data

The scRNA-seq output data were processed with the Cell Ranger pipeline (10×Genomics) against the mouse reference (mm10) datasets. Te gene-barcode matrices were analyzed and visualized using the Seurat R package (version 4.0.1)26. Genes expressed in less than 3 cells and cells containing less than 200 genes were filtered out. To remove potential doublets and low-quality cells, cells with unique gene counts of more than 6500 or a mitochondrial gene percentage greater than 10% were discarded.

### Flow cytometer analysis

Tongue epithelial cells were isolated as the method of preparation of TBOs. Mouse jejunum intestinal tissues were treated with 5mM EDTA for 30 min at 4□ and then shaken to fall down CRYPTs and villus. The isolated CRYPTs and villus were separated by filtration with 70 μm cell strainer. The flow through fraction mainly contains CRYPTS and the filtered out fraction mainly contains villus. Each fraction was treated with TrypLE express, and dissociated cells were used as intestinal epithelial cells. TBOs were digested with TrypLE express. The dissociated tongue epithelial, intestinal epithelial, and TBO cells were fixed with 4% paraformaldehyde solution for 15 min at RT. The fixed cells were permeabilized with 0.1% Triton-X100 and stained with primary antibodies and secondary antibodies. Mouse uterus were cut in pieces and treated with dispase on ice for 30 min, RT for 10 min, and 37 □ for 5 min. Then, the uterine epithelium was peeled off by pipetting. The uterine epithelial sheets were digested with TrypLE express. The dissociated cells were directly stained with fluorescent tagged antibodies against surface marker proteins. Stained cells were analyzed by SH800S cell sorter.

### Mouse taste tests

Mice were housed in two isolators whose light/dark cycles have a 12 h time difference. Those mice were subjected to two water bottles; one is pure water and the other is water with tastants at ZT0 or ZT12 for 6 hrs. The position of bottles were exchanged after 3 hrs to exclude positional effects. The consumption of water was measured by the weight.

## Author contributions

T.M. and K.M. performed all experiments. T.M. designed and supervised the experiments, and wrote the manuscript.

## Acknowledgments

We thank Genome Information Research Center, Research Institute for Microbial Diseases, Osaka University, Osaka, Japan for scRNAseq analysis. This work was supported by a grant from the Ministry of Education, Culture, Sports, Science and Technology of Japan to T. M. (19K08454) and the KMU consortium fund from Kansai Medical University.

## References

Aihara, E., Mahe, M.M., Schumacher, M.A., Matthis, A.L., Feng, R., Ren, W., Noah, T.K., Matsu-ura, T., Moore, S.R., Hong, C.I., et al. (2015). Characterization of stem/progenitor cell cycle using murine circumvallate papilla taste bud organoid. Sci Rep 5, 17185.

Bao, L., Roediger, J., Park, S., Fu, L., Shi, B., Cheng, S.Y., and Shi, Y.B. (2019). Thyroid Hormone Receptor Alpha Mutations Lead to Epithelial Defects in the Adult Intestine in a Mouse Model of Resistance to Thyroid Hormone. Thyroid 29, 439–448.

Bao, L., Shi, B., and Shi, Y.B. (2020). Intestinal homeostasis: a communication between life and death. Cell Biosci 10, 66.

Bjarnason, G.A., Jordan, R.C., and Sothern, R.B. (1999). Circadian variation in the expression of cell-cycle proteins in human oral epithelium. Am J Pathol 154, 613–622.

Chatterjee, A., and Hardin, P.E. (2010). Time to taste: circadian clock function in the Drosophila gustatory system. Fly (Austin) 4, 283–287.

Dorr, W., and Kummermehr, J. (1991). Proliferation kinetics of mouse tongue epithelium under normal conditions and following single dose irradiation. Virchows Arch B Cell Pathol Incl Mol Pathol 60, 287–294.

Fujimura, A., Kajiyama, H., Tateishi, T., and Ebihara, A. (1990). Circadian rhythm in recognition threshold of salt taste in healthy subjects. Am J Physiol 259, R931–935.

Geyfman, M., Kumar, V., Liu, Q., Ruiz, R., Gordon, W., Espitia, F., Cam, E., Millar, S.E., Smyth, P., Ihler, A., et al. (2012). Brain and muscle Arnt-like protein-1 (BMAL1) controls circadian cell proliferation and susceptibility to UVB-induced DNA damage in the epidermis. Proc Natl Acad Sci U S A 109, 11758–11763.

Grechez-Cassiau, A., Rayet, B., Guillaumond, F., Teboul, M., and Delaunay, F. (2008). The circadian clock component BMAL1 is a critical regulator of p21WAF1/CIP1 expression and hepatocyte proliferation. J Biol Chem 283, 4535–4542.

Guzman-Marin, R., Suntsova, N., Bashir, T., Szymusiak, R., and McGinty, D. (2007). Cell proliferation in the dentate gyrus of the adult rat fluctuates with the light-dark cycle. Neurosci Lett 422, 198–201.

Hao, Y., Hao, S., Andersen-Nissen, E., Mauck, W.M., 3rd, Zheng, S., Butler, A., Lee, M.J., Wilk, A.J., Darby, C., Zager, M., et al. (2021). Integrated analysis of multimodal single-cell data. Cell 184, 3573–3587 e3529.

Hofer, D., and Drenckhahn, D. (1996). Cytoskeletal markers allowing discrimination between brush cells and other epithelial cells of the gut including enteroendocrine cells. Histochem Cell Biol 105, 405–412.

Hong, C.I., Zamborszky, J., Baek, M., Labiscsak, L., Ju, K., Lee, H., Larrondo, L.F., Goity, A., Chong, H.S., Belden, W.J., et al. (2014). Circadian rhythms synchronize mitosis in Nemospora crassa. Proc Natl Acad Sci U S A 111, 1397–1402.

Huang, S., Kuri, P., Aubert, Y., Brewster, M., Li, N., Farrelly, O., Rice, G., Bae, H., Prouty, S., Dentchev, T., et al. (2021). Lgr6 marks epidermal stem cells with a nerve-dependent role in wound re-epithelialization. Cell Stem Cell 28, 1582–1596 e1586.

Kim, Y.S., and Ho, S.B. (2010). Intestinal goblet cells and mucins in health and disease: recent insights and progress. Curr Gastroenterol Rep 12, 319–330.

Kowalska, E., Ripperger, J.A., Hoegger, D.C., Bruegger, P., Buch, T., Birchler, T., Mueller, A., Albrecht, U., Contaldo, C., and Brown, S.A. (2013). NONO couples the circadian clock to the cell cycle. Proc Natl Acad Sci U S A 110, 1592–1599.

Matsu-Ura, T., Dovzhenok, A., Aihara, E., Rood, J., Le, H., Ren, Y., Rosselot, A.E., Zhang, T., Lee, C., Obrietan, K., et al. (2016). Intercellular Coupling of the Cell Cycle and Circadian Clock in Adult Stem Cell Culture. Mol Cell 64, 900–912.

Matsuo, T., Yamaguchi, S., Mitsui, S., Emi, A., Shimoda, F., and Okamura, H. (2003). Control mechanism of the circadian clock for timing of cell division in vivo. Science 302. 255–259.

Mendez-Ferrer, S., Lucas, D., Battista, M., and Frenette, P.S. (2008). Haematopoietic stem cell release is regulated by circadian oscillations. Nature 452, 442–447.

Moses, R., and Kummermehr, J. (1986). Radiation response of the mouse tongue epithelium. Br J Cancer Suppl 7, 12–15.

Nakamura, Y., Sanematsu, K., Ohta, R., Shirosaki, S., Koyano, K., Nonaka, K., Shigemura, N., and Ninomiya, Y. (2008). Diurnal variation of human sweet taste recognition thresholds is correlated with plasma leptin levels. Diabetes 57, 2661–2665.

Nikhil, K.L., Korge, S., and Kramer, A. (2020). Heritable gene expression variability and stochasticity govern clonal heterogeneity in circadian period. PLoS Biol 18, e3000792.

Plikus, M.V., Vollmers, C., de la Cruz, D., Chaix, A., Ramos, R., Panda, S., and Chuong, C.M. (2013). Local circadian clock gates cell cycle progression of transient amplifying cells during regenerative hair cycling. Proc Natl Acad Sci U S A 110, e2106-2115.

Smaaland, R., Sothern, R.B., Laerum, O.D., and Abrahamsen, J.F. (2002). Rhythms in human bone marrow and blood cells. Chronobiol Int 19, 101–127.

Stuart, T., Butler, A., Hoffman, P., Hafemeister, C., Papalexi, E., Mauck, W.M., 3rd, Hao, Y., Stoeckius, M., Smibert, P., and Satija, R. (2019). Comprehensive Integration of Single-Cell Data. Cell 177, 1888–4902 e1821.

Umemura, Y., Koike, N., Ohashi, M., Tsuchiya, Y., Meng, Q.J., Minami, Y., Hara, M., Hisatomi, M., and Yagita, K. (2017). Involvement of posttranscriptional regulation of Clock in the emergence of circadian clock oscillation during mouse development. Proc Natl Acad Sci U S A 114, e7479–E7488.

Wysokinski, A., Sobow, T., Kloszewska, I., and Kostka, T. (2015). Mechanisms of the anorexia of aging-a review. Age (Dordr) 37, 9821.

Yagita, K., Horie, K., Koinuma, S., Nakamura, W., Yamanaka, I., Urasaki, A., Shigeyoshi, Y., Kawakami, K., Shimada, S., Takeda, J., et al. (2010). Development of the circadian oscillator during differentiation of mouse embryonic stem cells in vitro. Proc Natl Acad Sci U S A 107, 3846–3851.

Yang, Q., Pando, B.F., Dong, G., Golden, S.S., and van Oudenaarden, A. (2010). Circadian gating of the cell cycle revealed in single cyanobacterial cells. Science 327, 1522–1526.

Zhang, Z.B., Sinha, J., Bahrami-Nejad, Z., and Teruel, M.N. (2022). The circadian clock mediates daily bursts of cell differentiation by periodically restricting cell-differentiation commitment. Proc Natl Acad Sci U S A 119, e2204470119.

