## Supplementary figure legends for "Circadian clock-gated cell renewal controls time-dependent changes in taste sensitivity"

***Supplemental figure legends***

**Figure S1 *Time course expression of stem/progenitor and differentiated cell marker genes in TBOs*.** (**a**) mRNA expression of core clock genes (*Bmal1* and *Per2*), adult stem cell marker genes, (*Lgr6*, *Axin2* and *Bmi1*), and differentiated taste cell marker genes (*Krt8*, *α-Gustducin* (*Gnat3*), *NTPDase2*, and *T1R2*) in *Bmi1* knock-down TBOs. (**b**) mRNA expression of taste marker genes.

**Figure S2 *Cell type annotations of scRNAseq and time-dependent changes of cell types in tongue epithelium*.** (**a**-**c**) Annotations of types of cells are shown by the gene expression patterns. Detection of stem/progenitor cell marker genes (**a**), progenitor cell marker genes (**b**), and differentiated cell marker genes (**c**) in the tongue epithelium. (**d**-**e**) Flow cytometry experiments of pan-taste cell marker (ENTPD2) or type II taste cell marker (GNAT3) expressing cells in TBOs (**d**) and tongue epithelium (**e**) at different circadian time points.

**Figure S3 *Heterogeneity of clock gene expressions in tongue epithelium*.** (**a**) mRNA expression of a taste stem cell marker *Lgr6* and taste differentiated cell markers, *Gnat3*, *NTPDase2*, and *K8* in TBOs cultured in control and stem cell rich conditions. (**b**-**d**) Expression maps of circadian morning genes (**b**), night genes (**c**), and the superimposed map of those circadian clock genes (**d**).

**Figure S4 *Time-dependent changes of apoptotic cells and cellular populations in the intestine and the uterus*.** (**a**) Apoptotic cells were detected by active CASPASE3 antibody, and tuft cells were detected by KRT18 antibody in intestinal CRYPT and villus epithelium. (**b**) Apoptotic cells were detected by ANNEXIN V antibody, and stem cells were detected by CD146 antibody in uterine epithelium.

**Figure S5** ***Salty and sour taste sensing in tongue epithelium at different time points.*** Mean taste preference ratios of ZT0 and ZT12 mice using 6-hour two-bottle (tastant versus distilled water) preference tests. Behavioral responses of WT (upper panels) and *Lgr6^CreERT/+^/R26^DTA/+^* mice (lower panels) to salty (**a**, NaCl: 50, 100, 200, and 300 mM) and sour (**b**, HCl: 0.1, 1, 3, and 10 mM) tastants. The values and errors correspond to the average values and the SDs, respectively. **p* < 0.05, student’s *t*-test.
