## Supplementary material for "Circadian clock-gated cell renewal controls time-dependent changes in taste sensitivity": Table S1

| Taste cell type | Marker genes |
|---|---|
| Type I | *Entpd2* , *Slc1a3* |
| Type II | *Tas1r1* , *Tas1r2* , *Gnat3* , *Tas2r102* , *Tas2r104* , *Tas2r105* , Tas2r107, *Tas2r108* , *Tas2r110* , *Tas2r113* , *Tas2r117* , *Tas2r120* , *Tas2r135* , *Tas2r137* , *Tas2r138* |
| Type III | *Car4* , *Pkd2l1* , *Ddc* , *Scnn1a* , *Scnn1b* , *Scnn1g* , *Snap25* |
| Pan | *Krt8* , *Kcnq1* |

| Number of expressing cells in ZT0 | Number of expressing cells in ZT12 |
|---|---|
| 862 (1405) | 1349 (2352) |
| 35 (1405) | 25 (2352) |
| 700 (1405) | 1193 (2352) |
| 36 (1405) | 51 (2352) |

* Total cell numbers are in parencesis.
