## Supplementary figures and images for "Circadian clock-gated cell renewal controls time-dependent changes in taste sensitivity"

### Figure S1

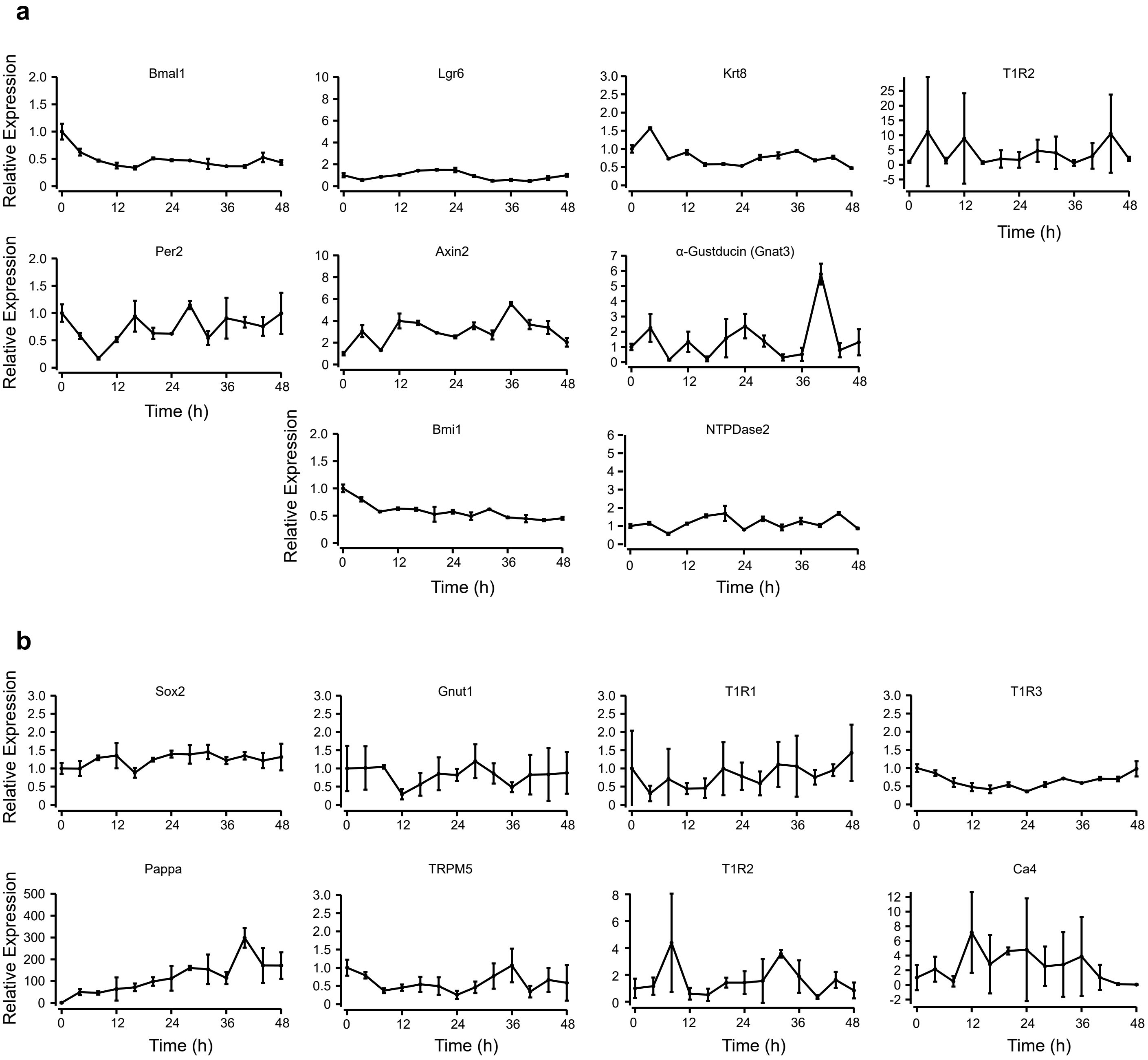

### Figure S2

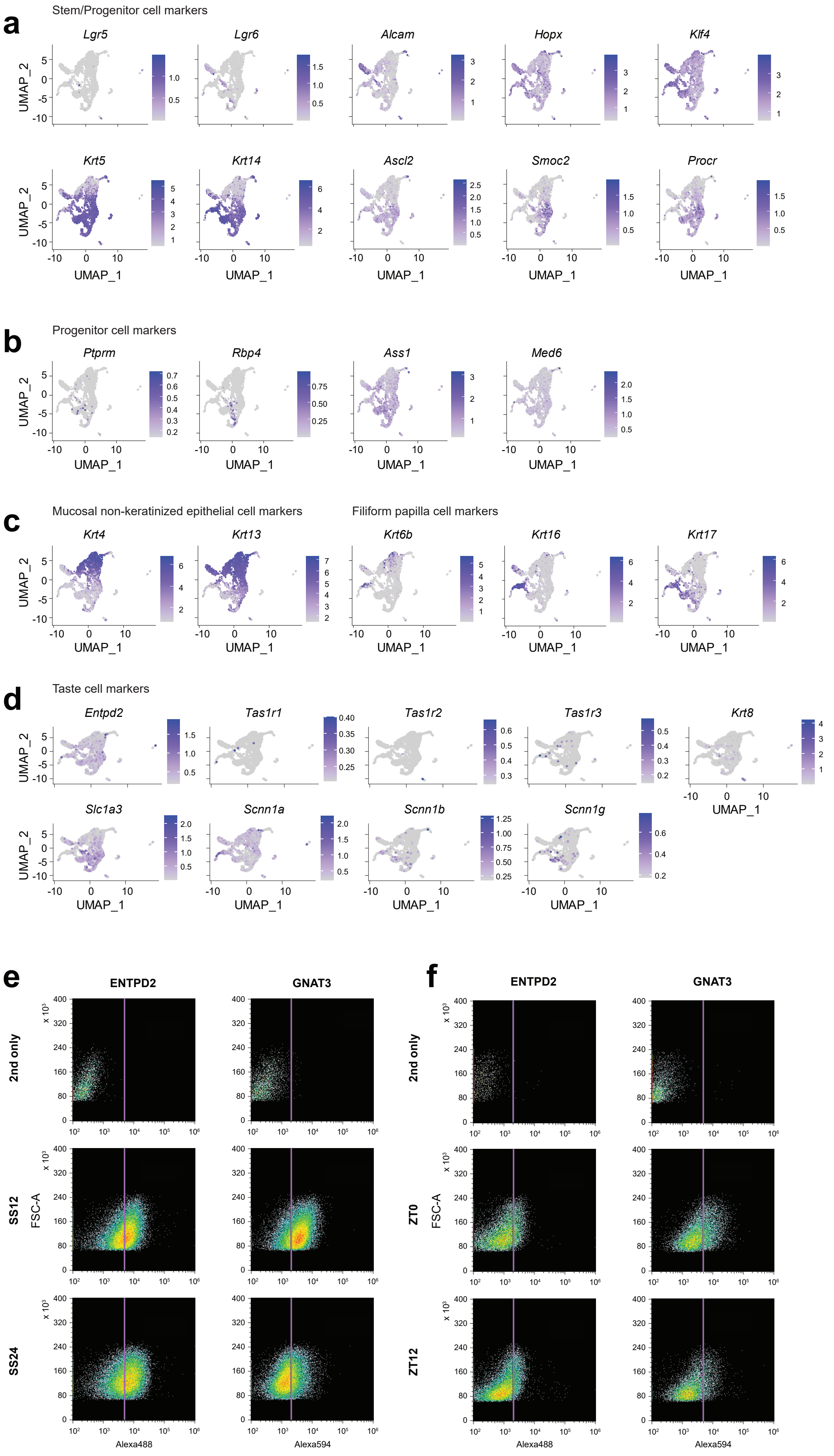

### Figure S3

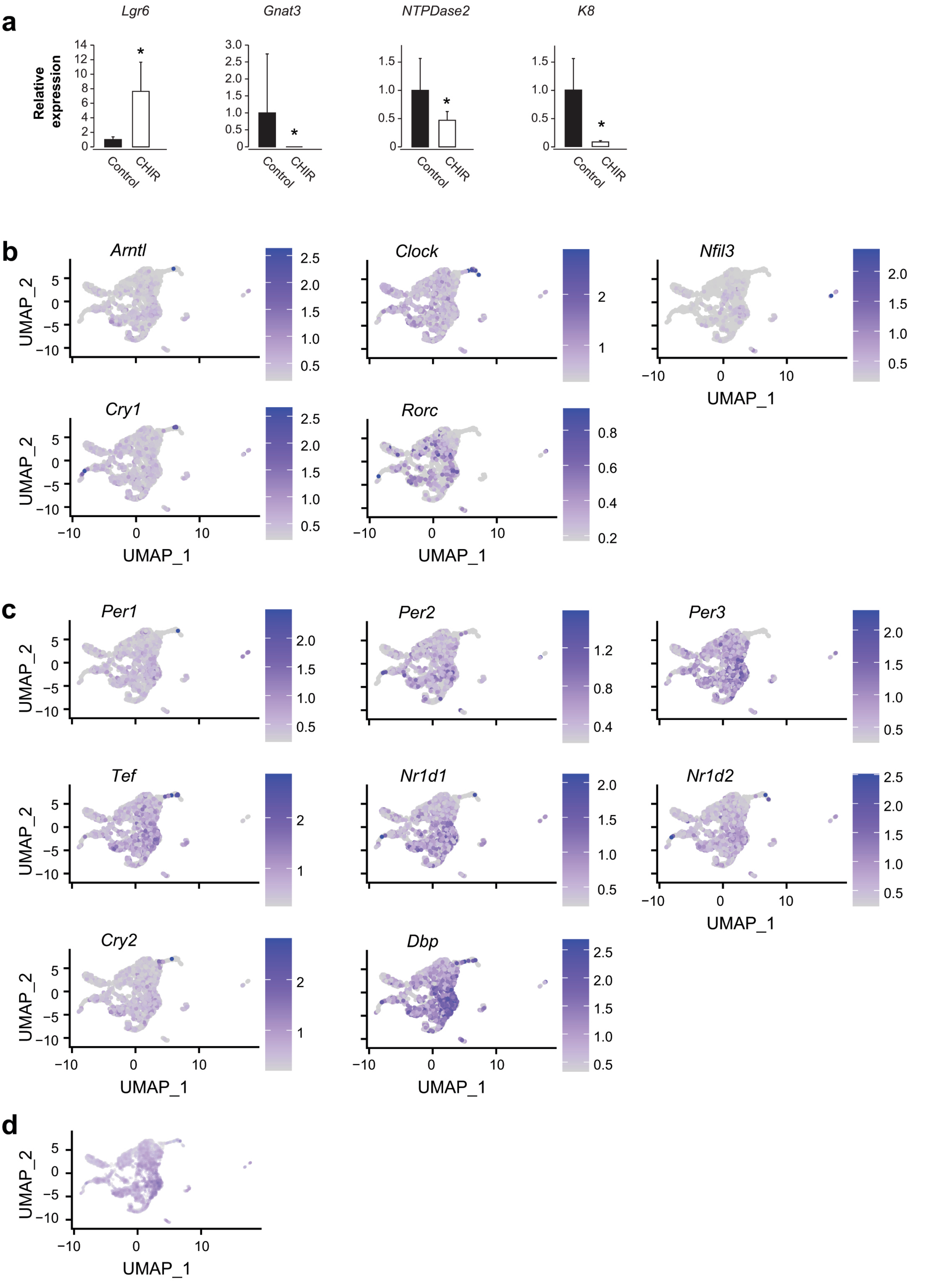

### Figure S4

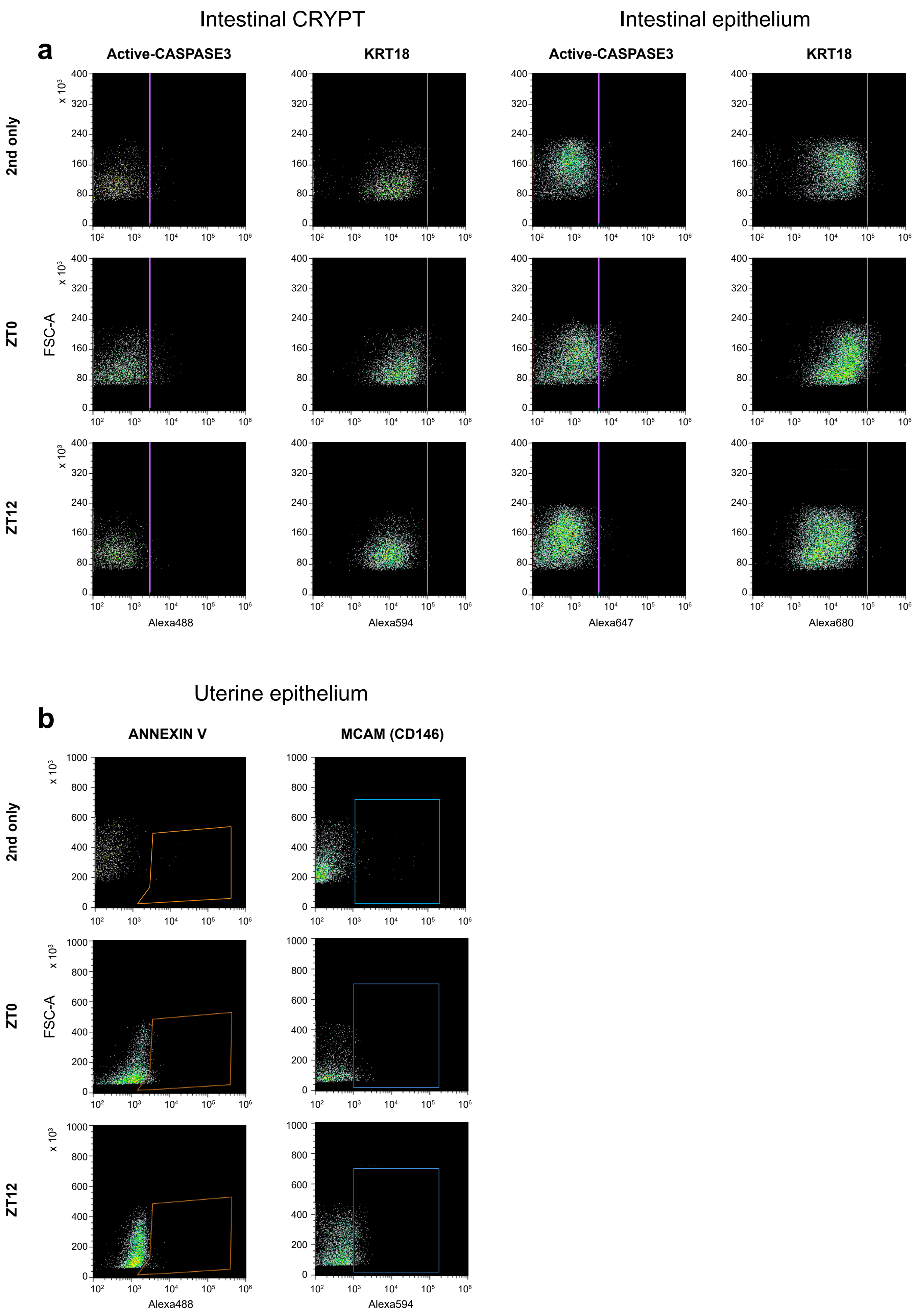

### Figure S5

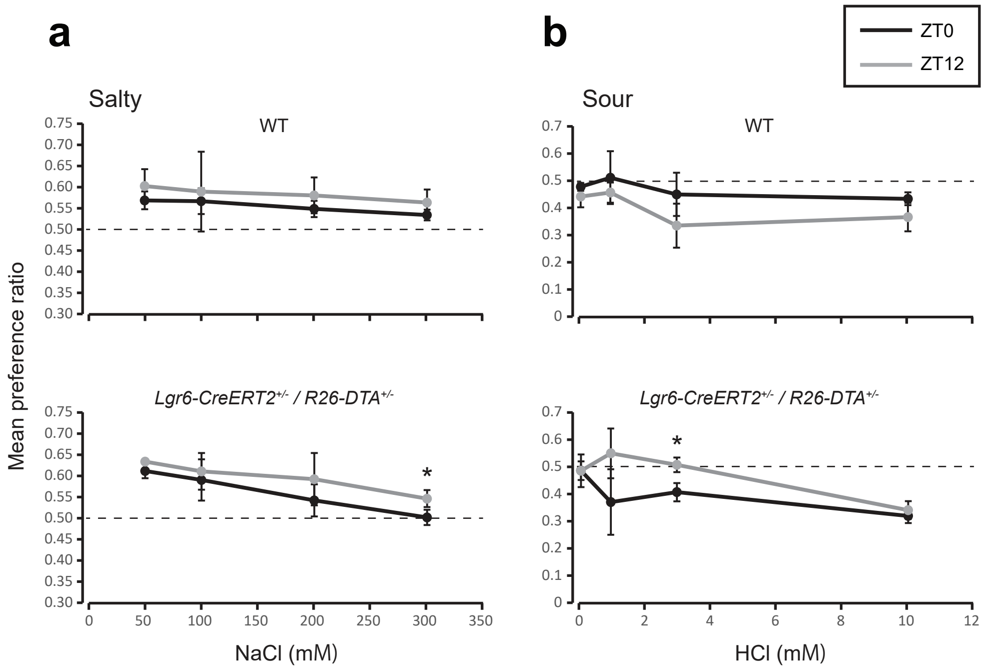
